# Adaptation in isolated populations: when does it happen and when can we tell?

**DOI:** 10.1101/052050

**Authors:** Jessica L. Crisci, Matthew D. Dean, Peter Ralph

## Abstract

Isolated populations with novel phenotypes present an exciting opportunity to uncover the genetic basis of ecologically significant adaptation, and genomic scans for positive selection in such populations have often, but not always, led to candidate genes directly related to an adaptive phenotype. However, in many cases these populations were established by a severe bottleneck, which can make identifying targets of selection problematic. Here we simulate severe bottlenecks and subsequent selection on standing variation, mimicking adaptation after establishment of a new small population, such as an island or an artificial selection experiment. Using simulations of single loci under positive selection and population genetics theory, we examine how population size and age of the population isolate affects the ability of outlier scans for selection to identify adaptive alleles using both single site measures and haplotype structure. We find and explain an optimal combination of selection strength, starting frequency, and age of the adaptive allele, which we refer to as a Goldilocks zone, where adaptation is likely to occur, and yet the adaptive variants are most likely to derive from a single ancestor (a “hard” selective sweep); in this zone, four commonly used statistics detect selection with high power. Real-world examples of both island colonization and experimental evolution studies are discussed. Our study provides concrete considerations to be made before embarking on whole genome sequencing of differentiated populations.

## Introduction

Populations that have colonized islands or become demographically isolated in new habitats can be subject to strong selection as they adapt to their new environment (Losos & Rickfels; Pergams & Lawler 2009). This often leads to the evolution of new phenotypes, making isolated populations a useful model to study evolution in action (e.g. Losos *et al.* 2001; Gill 1977; Reznick & Bryga 1987), especially if mainland or source populations can serve as an ancestral comparison. A number of recently established examples come from human introduction of small rodents on islands (Berry 1996; Martinkova *et al*. 2013; Pergams *et al.* 2015; Patton *et al*. 1975; Ledevin *et al*. 2016), providing a “natural experiment”.

However, researchers are faced with a troublesome problem: it is quickly becoming easier and less expensive to sequence whole genomes of many individuals, but with the mixed success of genomes scans for identifying selection (reviewed in Akey 2009) the utility of such genome-first approaches in uncovering genetic causes of adaptive phenotypes is unclear.

Among the top concerns is the ability to detect sites affected by selection given the often severe bottlenecks that accompany colonization events (Keller & Taylor 2008). For instance, Poh *et al*. (2014) were unable to retrieve significant signal of positive selection from the region surrounding a well-characterized adaptive mutation for light coat color in a population of Florida beach mice. However, they did successfully identify a significant signal at a similarly well-characterized adaptive mutation in a population of Nebraska Sand Hills mice, perhaps because the much larger ancestral effective population size of this second population (*N*_e_ of 50,000 vs. *N*_e_ of 2,500 in the beach mice) allowed the signal to persist for much longer.

Experimental evolution studies also involve local adaptation of isolated populations to novel environmental challenges, and so many experimental designs are covered by the same discussion. A diverse literature has documented genetically based phenotypic divergence as a result of many different selective regimes (e.g. Wright & Dobzhansky 1945; Reznick & Endler 1982; Holland & Rice 1999; Firman & Simmons 2011; Kolbe *et al.* 2012).

When a single copy of a new beneficial allele appears in a population it can rise (“sweep”) rapidly to fixation. Such classic or “hard” selective sweeps reduce genetic variation and increase haplotype lengths around the selected site because the time to the most recent common ancestor is shorter at that site (Fisher 1918, Maynard Smith & Haigh 1974, Kaplan *et al*. 1989). However, in populations that have recently experienced a bottleneck, selection has a higher chance of success if it acts on sites already segregating in the population, especially in scenarios when selection is weak (Hermisson & Penninngs 2005). In such “soft” sweeps, a beneficial allele that is already present on multiple genetic backgrounds goes to fixation, which does not necessarily result in loss of linked variation. The segregating allele could have one origin, i.e., all haplotypes carrying the allele are identical by descent, or it could have come from multiple de novo mutations. Either way, this reduces the footprint of selection around the target site, owing to the fixation of haplotypes from multiple genetic backgrounds (Przeworski *et al*. 2005).

To determine how much power we have to detect selected sites from whole genome population sequence we simulate selection on genetic variation under several demographic scenarios that vary the timing and strength of a bottleneck, the selection coefficient, and the starting frequency of the selected variant. We ask two main questions: 1) Under what scenarios will selection overcome drift?, and 2) When selection overcomes drift, can we detect it? We test for selection using four general statistics: F_ST_ (Wright 1950), OmegaPlus (Alachiotis *et al*. 2012), H12 (Garud *et al*. 2015) and nucleotide diversity (π).

To detect a region under selection using a given test statistic, selection must significantly distort the genealogies near the selected site so as to move the value of the statistic, and this distortion must be distinguishable from the effects of neutral demography. In a small, isolated population undergoing adaptation from standing variation, the key descriptors are the number of copies of the selected allele that successfully sweep; the speed of the sweep; and the time since selection began.

We find that power is maximized at intermediate values of the key parameters, a “Goldilocks zone” in parameter space: all four statistics applied here perform the best if initial frequency is low (but not too low), selection is strong, and the population is not too old and not too young. (The name comes from Goldilocks’ fabled preference for intermediate chair size, porridge temperature, and bed softness.) We apply results to real world examples, finding that detecting selection may be difficult in many vertebrate systems, or at least require the strength of selection, initial population diversity, and size and age of the population to fall within particular bounds.

## Methods

### Msms Simulations

We used the program msms (Ewing & Hermisson 2010) to simulate genetic variation after adaptation in the following situations. An ancestral population of effective site *N*_*e*_ splits *t* generations ago into two populations—one population is bottlenecked to a smaller size *N*_*I*_=*nN*_*e*_ (and remains this size subsequently), and the other remains at the ancestral population size (Figure 2). The bottlenecked population experiences selection from the time of the split to the time of sampling. We simulated 500 replicates of a 1Mb region, sampling 50 diploids (100 chromosomes) each replicate. The ancestral population size, *N*_e_, was set to 70,000, and the bottleneck event occurred at either *t* = 50, 250, or 1000 generations in the past. We simulate three different selection coefficients as follows: *s*_AA_ = 0.001 *s*_AA_ = 0.01; or *s*_AA_ = 0.1, where “A” is the derived, beneficial allele and “a” is the ancestral allele. In each case selection is additive; *s*_Aa_*=s*_AA_/2. The starting frequency of the beneficial allele was either 0.01, to more closely resemble a hard sweep, or 0.1 to represent a soft sweep. The ratio of the new bottlenecked population size to the ancestral population size, *n*, was set to 0.001, 0.01, or 0.02, which creates isolated populations with sizes, *N*_I_, equal to 70, 700, and 1400 individuals, respectively. All combinations of these parameters yield 54 models experiencing selection, as well as 9 neutral controls (one for each combination of bottleneck size and age of the population).

**Fig. 2.**
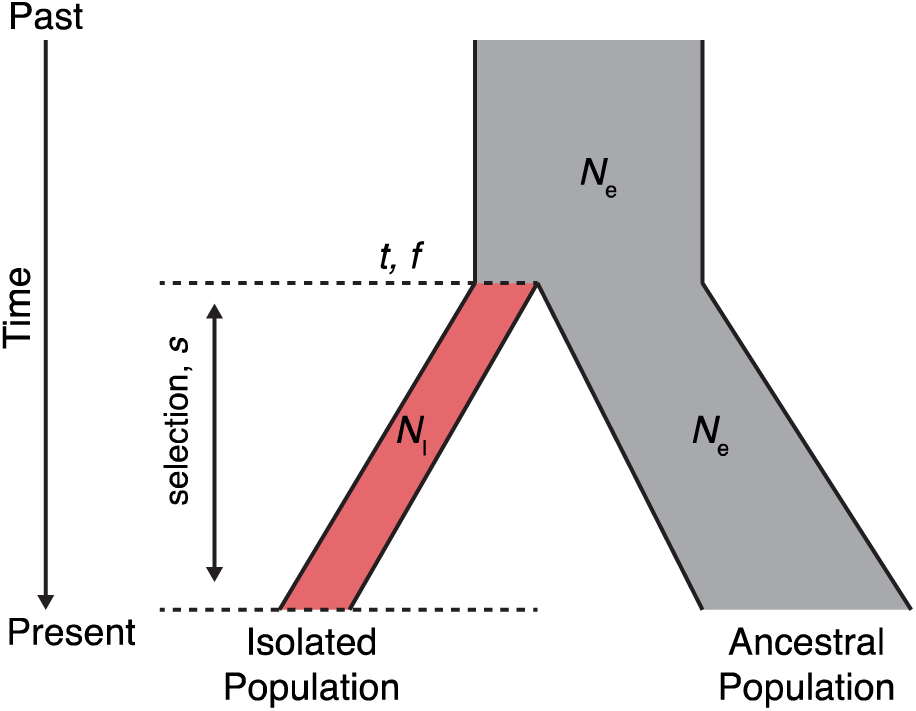
Summary of the model used to simulate isolated populations. Both populations remain at a constant size. The isolated population is formed by a proportion, *n*, of the ancestral population, where *nN*_e_ = *N*_I_. Selection begins simultaneously with the bottleneck at time *t* generations in the past on an allele at starting frequency, *ƒ*, in the isolated population. Note that we don’t condition on fixation of the beneficial allele.

### Statistical tests for detecting selection

We applied four statistics to detect selection from our simulations. First, we examined genetic differentiation between populations, quantified by *F*_ST_, since selection in the bottlenecked population should increase its genetic divergence to the ancestral population in the region of the selected mutation. Second, we calculated mean pairwise nucleotide divergence per bp, quantified by π, since selection should decrease variation in the region of the selected mutation. We expect these statistics to be somewhat complimentary since *F*_ST_ is sensitive to differentiation between allele frequencies in two populations, whereas π is only concerned with the number of differences between alleles in a single population. We obtained π using the-oTPi flag in msms and compute *F*_ST_ using allele frequencies directly from the msms output, with *F*_*ST*_=(*H*_*T*_ -*H*_*S*_)/*H*_*T*_, where *H*_T_ is the expected total heterozygosity in both populations (the probability that two randomly chosen alleles from the entire sample differ), and *H*_S_ is the expected heterozygosity in each subpopulation (the probability that two randomly chosen alleles from the same subpopulation differ).

We used two additional statistics based on the signature of linkage disequilibrium (LD). LD is expected to be high on either side of a selected site due to linked neutral variation and reduced across the site (McVean 2007). OmegaPlus (Alachiotis *et al.* 2012)is a sliding window approach to detect this pattern based on Kim & Nielsen’s (2004) ω_MAX_. We also consider H12, a statistic designed to be sensitive to sweeps from standing variation. H12 quantifies haplotype homozygosity after combining the two most frequent haplotypes into one class so that **H12** = 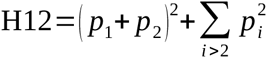 where *p* _i_ is the frequency of the *i*^th^ haplotype in a window of fixed size (Garud *et al.* 2015). Both OmegaPlus and H12 use the signals left behind by recombination during the sweep to identify targets of selection, but we do not expect extensive overlap between the two statistics: OmegaPlus uses correlations between individual alleles within a window to find specific patterns of LD decay while H12 is concerned with haplotype structure.

### Determining power of statistics

To determine power to detect selection, we first had to identify the parameter space over which selection can be effective. We removed from further analysis any replicates where the final frequency of the beneficial allele was less than 0.5 or there were fewer than 100 segregating sites across the 1 Mb region. Even though we removed individual replicates from each model, some models were completely eliminated for not having any replicates that passed either filter. The segregating sites filter removed many models with the most severe bottleneck (*N* _I_ = 70), where drift in the extremely small population caused fixation of most variation (see below). Also, many models from the youngest bottleneck (*t* = 50 generations) did not contain any replicates where the frequency of the beneficial allele made it above 0.5.

For the remaining replicates we computed each of the 4 statistics. F_ST_ and π were calculated in non-overlapping windows of 100Kb. H12 was calculated in windows of 100 SNPs with a step size of 10. OmegaPlus was calculated with maxWin set to 150Kb and minWin set to 10Kb. We looked at several window sizes for each statistic and chose the sizes that were best able to detect the signal of the selective sweep.

We then determined the 99^th^ (F_ST_, OmegaPlus, H12) or 1^st^ (π) percentile value for each neutral demographic scenario across all windows from all 500 replicates and used this as a threshold value for the corresponding demographic models with selection. To determine power, we calculated the proportion of replicates under each model whose maximum window value was greater (or less in the case of π) than the threshold value. Power was not assessed in models that contained less than 20 replicates (4% of the 500 replicates) after filtering for final frequency and segregating sites.

### Application to real-world examples

We apply lessons from the simulations to island colonization events, as well as experimental evolution studies. Firman and Simmons (2008) discovered an example of phenotypically divergent mating ecologies among island mice off the coast of Western Australia, finding that the extent of multiple paternity varied across seven island populations and that males from populations with high multiple paternity developed larger testis and produced more sperm of higher quality—all traits which could indicate varied levels of sperm competition amongst males (Firman *et al*. 2013). Uncovering the underlying genetic causes of these diverse phenotypes would provide insight into an interesting ecological observation.

Two of their populations, Rat Island and Whitlock Island, showed high and low multiple paternity, respectively. We examined these populations in the context of the models that we simulate here, in order to assess whether they may be candidates for identifying selected sites. Census population sizes based on trapping data were estimated to be 772 and 111 individuals on Rat Island and Whitlock Island, respectively (Firman & Simmons 2008). Whitlock Island was likely initially colonized by shipwrecks as early as the 1600s. Rat Island was more likely colonized through its intermittent inhabitance by humans for the fishing and guano industries, starting in the mid 1800s. If we assume 2 generations per year for *M. domesticus* in the wild, we estimate that Rat Island was colonized around 300 generations ago and Whitlock around 800 generations ago.

We also consider another phenotypically interesting population of mice found on Gough Island, in the South Atlantic. These mice are extremely large and exhibit carnivorous behavior, feeding on the small seabird chicks on the island (Rowe-Rowe & Crafford 1992, Cuthbert and Hilton 2003). Grey *et al*. (2014) estimated a colonization event at around 110 generations ago and a founding population size of 950 individuals.

Below, we discuss the theoretical framework for a selective sweep and explore how it applies to populations that have suffered a recent bottleneck. Using the colonization time and population size estimates from real life island examples, we determine whether a beneficial allele could have survived the bottleneck event and if the allele could have had sufficient time to reach fixation in these populations given the selection scenarios represented by our simulations.

In addition, we compiled several examples typical of island adaptation and experimental evolution studies (Table 1), focusing on vertebrate systems, that usually most closely match the parameters we simulated under in this study (although the theoretical results should apply more generally). The parameters listed are taken from the literature cited when possible, and are meant to serve as illustrative examples rather than exact estimates for particular systems. Furthermore, the isolated population in our simulations remained at the same small size; in many real situations the population expands substantially after introduction (Reznick & Ghalambour 2001). Because of this, Table 1 has separate columns for “effective introduction size” and “long term effective size”. The former is used in calculations of the initial available diversity (*K* below), and the latter gives the time scale on which drift erases the initial signal (the upper boundary of the Goldilocks zone in Figure 1).

**Fig. 1.**
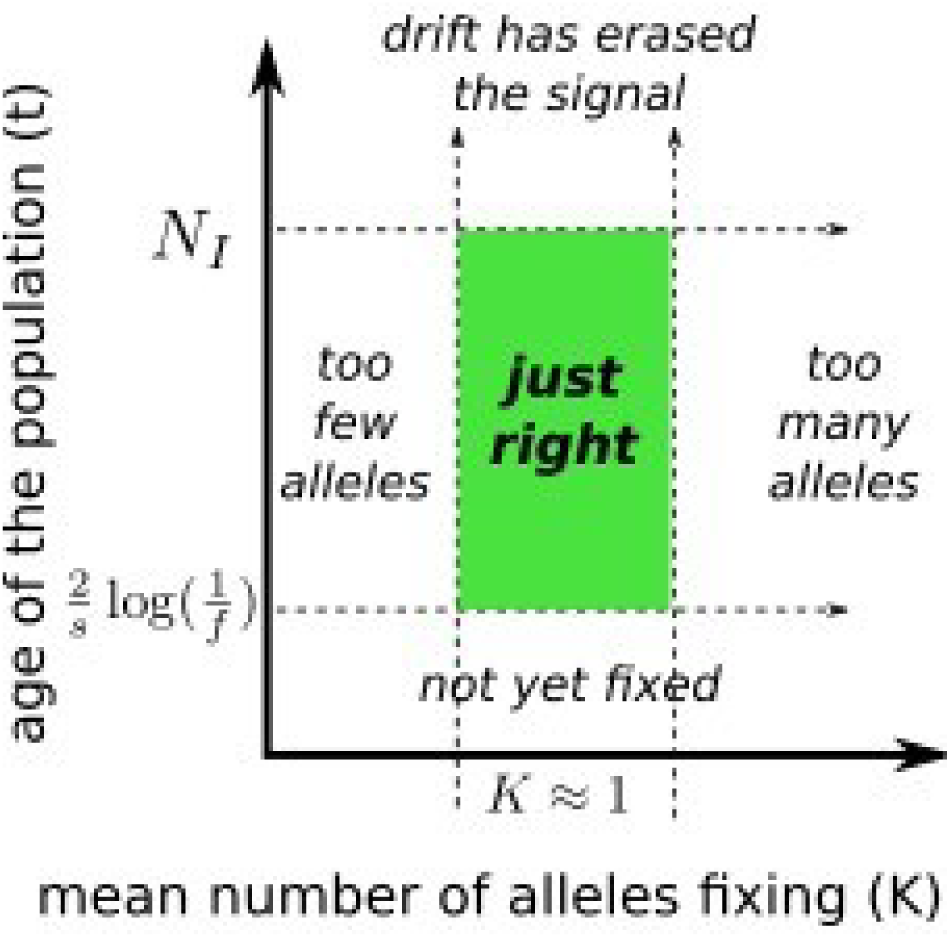
Conceptual diagram of the Goldilocks zone. In a recently formed, isolated population that has suffered a bottleneck, detecting selection depends on both having enough alleles to fix without fixing too many haplotypes – creating a sweep that is too soft to detect – and being able to observe the sweep before the signal is erased by drift. For this to be true, the number of generations since the beneficial allele arose, *t* must be longer than the time to fixation, and less than the effective population size, *N*_I_, the time scale of coalescence in the population. For selection to occur, the number of alleles expected to fix, *K* = 2*s*_*Aa*_*N*_I_*ƒ*, should be at least 1.

**Table 1.** Examples of adaptation in recently isolated populations. Parameters are taken from the cited papers, where possible; additionally, guppy (*Poecilia reticulata*) population size and age estimates are from Fraser *et al.* (2015). Both effective introduction size and long-term effective size correspond to *N*_I_ in the text. Derived parameters are computed as described in the text, and a scenario is “Goldilocks” if K is between 0.25 and 3, and if *t* is between and the long-term effective population size.

| Example | Founding $N_e$ | Long-term $N_e$ | Age (T, gens) | Maximum Goldilocks frequency (f) | Minimum Goldilocks selection (s) | Initial frequency (f) | Selection (s) | Number of sweeping haplotypes (K) | Duration of the sweep (Tfix) | Goldilocks? | |
| --- | --- | --- | --- | --- | --- | --- | --- | --- | --- | --- | --- |
| Laboratory selection on mouse running preference (Swallow et al. 1999) | 20 | 20 | 10 | 0.36 | 0.07 | 0.05 | 0.05 | 0.1 | 46 | NO | too short and too few alleles |
|  |  |  |  |  |  | 0.05 | 0.25 | 0.5 | 4 | YES |  |
| Experimental guppy introduction to low-predation streams (Endler 1980) | 200 | 30 | 15 | 0.01 | 0.25 | 0.05 | 0.05 | 1 | 46 | NO | too short and too few alleles |
|  |  |  |  |  |  | 0.05 | 0.25 | 5 | 4 | NO | too many alleles |
| Naturally isolated guppy populations in low-predation streams (Endler 1980) | 40 | 4000 | 1000 | no limit | 1/Ne | 0.05 | 0.05 | 0.2 | 46 | NO |  |
|  |  |  |  |  |  | 0.05 | 0.25 | 1 | 4 | YES |  |
| Recent earthquake uplift isolating marine stickleback in new freshwater ponds (Lescak et al. 2015) | 200 | 200 | 50 | 0.09 | 0.03 | 0.05 | 0.05 | 1 | 46 | YES |  |
|  |  |  |  |  |  | 0.05 | 0.25 | 5 | 4 | NO | too many alleles |
| Long-term experimental evolution of *Drosophila* for life history traits Burke et al. 2010) | 1000 | 1000 | 200 | 0.06 | 0.01 | 0.001 | 0.05 | 0.1 | 124 | NO | too few alleles |
|  |  |  |  |  |  | 0.001 | 0.25 | 0.5 | 8 | YES |  |
| Selection on coat color in island *Peromyscus* populations of Florida (Poh et al. 2014) | 200 | 50000 | 7000 | no limit | 1/Ne | 0.01 | 0.05 | 0.2 | 78 | NO | too long and too few alleles |
|  |  |  |  |  |  | 0.01 | 0.25 | 1 | 5 | YES |  |

## Results & Discussion

### Conditions that favor fixation of the beneficial allele

There are many factors that influence whether a selected allele will fix in a population. Fixation will depend largely on the strength of selection, the starting frequency of the beneficial allele, and the amount of time since directional selection began. In order to be successful, a selected allele must escape loss due to demographic stochasticity and then have enough time to reach an appreciable frequency. First, recall that the probability that a beneficial allele fixes in a diploid population is approximately 2*s*_*Aa*_ (Haldane 1924) divided by the variance in haploid fitness, which we take to be 1. In a bottlenecked population, initially there are *N*_I_*ƒ* beneficial alleles present, of which

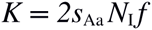

are expected to escape drift and potentially reach fixation (shown in Table 2). These alleles will be located on unique haplotypes if each was present on a different genetic background prior to the bottleneck event, in which case *K* is the expected number of haplotypes that sweep. The allele increases in frequency by roughly 1 + *s*_*Aa*_ every generation. If it escapes drift the frequency after *T* generations will be around *ƒ*_*T*_=*ƒ*(1+*s*_*Aa*_)^*T*^, and so the time, *T*, it takes to go from *ƒ* to *ƒ*’ is around

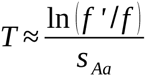

**Table 2.** **Estimated No. of Fixed Beneficial Alleles**

| Selection ($s$) | Starting Frequency ($f$) | | | | | |
| --- | --- | --- | --- | --- | --- | --- |
|  | 0.01 |  |  | 0.1 |  |  |
|  | 0.0005 | 0.005 | 0.05 | 0.0005 | 0.005 | 0.05 |
| $n = 0.001$ | 0.0007 | 0.007 | 0.07 | 0.007 | 0.07 | 0.7 |
| $n = 0.01$ | 0.007 | 0.07 | 0.7 | 0.07 | 0.7 | 7 |
| $n = 0.02$ | 0.014 | 0.14 | 1.4 | 0.14 | 1.4 | 14 |

The mean time to reach a frequency of *ƒ*’ = 0.5 for the *ƒ* and *s* combinations simulated here are shown in Table 3; the frequency 0.5 is certainly not “fixed”, but is chosen to be conservative.

**Table 3.** **No. of Generations to Reach f = 0.5**

| | Starting Frequency ($f$) | |
| --- | --- | --- |
|  | 0.01 | 0.1 |
| $s = 0.0005$ | 7824.0 | 3218.9 |
| $s = 0.005$ | 782.4 | 321.9 |
| $s = 0.05$ | 78.2 | 32.2 |

From these estimates we can see at the lower starting frequency, *ƒ* = 0.01, among our simulations only the strongest selection, *s*_*Aa*_ = 0.05, is expected to yield enough alleles with the potential to fix, and only for the less severe bottleneck parameters (Table 2, grey cells). Indeed, only these cases produce a high number of replicates that reach at least frequency 0.5, at all 3 time points (Figure 3A). We also see that these initially low frequency alleles take substantially more time to reach fixation when compared with the alleles from higher starting frequency, *ƒ* = 0.1 (Figure 4). When selection is very weak, *s*_*Aa*_ = 0.0005, few alleles escape drift in these small populations, as shown in Table 2 and the black lines in Figure 3. When the initial frequency is higher, *ƒ* = 0.1, fixation becomes possible for *s* = 0.005 for the two less severe bottleneck models. From Table 3 the time it would take for one of these alleles to reach frequency 0.5 is 321 generations, which explains why we only see a large proportion of simulations fixing at 1000 generations under this model (Fig 3B, red lines). When selection is strongest (*s*_*Aa*_ = 0.05) fixation happens very quickly, with the beneficial allele reaching frequency 0.5 in 32 generations from *ƒ* = 0.1, and 78 generations when *ƒ* = 0.01 (Table 3 and Figure 3, blue lines).

**Fig. 3.**
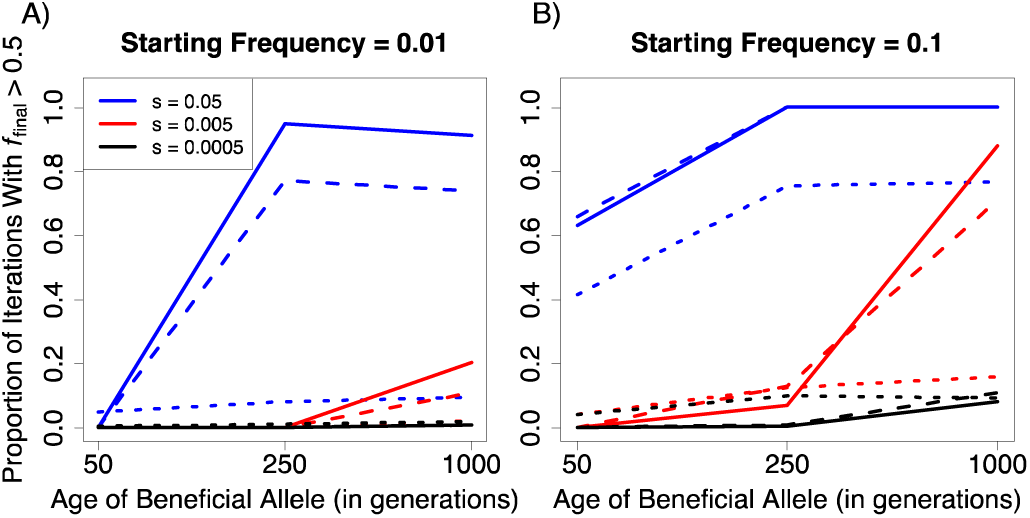
Proportion of simulations with final frequency of beneficial allele greater than 0.5. A total of 500 replicates were simulated for each set of parameters. Solid lines represent the models with the least severe bottleneck, where *N*_I_=1400; dashed lines, *N*_I_=700; dotted lines are the most severe bottleneck, *N*_I_=70.

**Fig. 4.**
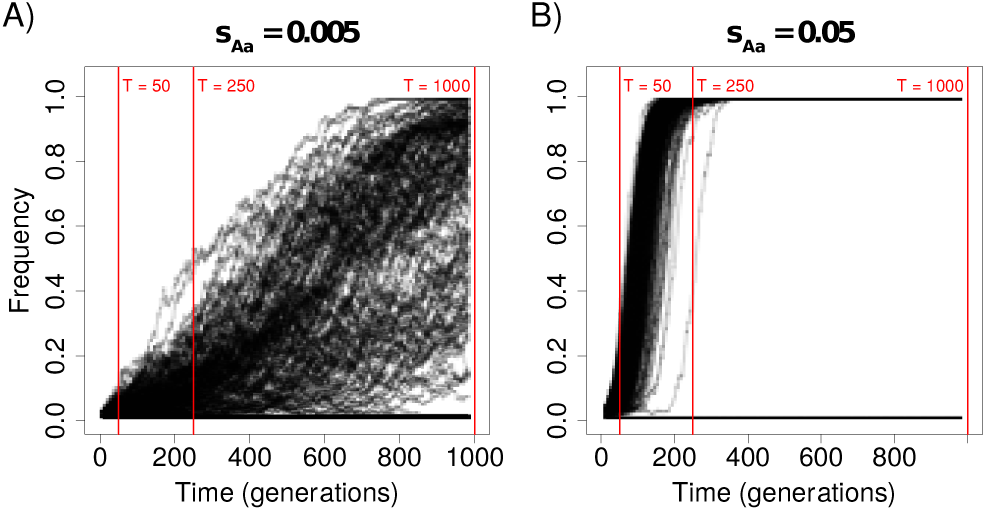
Trajectories from two selection coefficients. In each, the population size is *N*_I_ = 1400 and the starting frequency of the allele is *ƒ*=0.01. All 500 replicates are shown on each plot. A) The beneficial allele is under weak selection, *s*_Aa_ = 0.005, and has a reasonable probability of fixing, but the sweep is still ongoing at 1000 generations. B) The beneficial allele is under strong selection and fixes within 250 generations.

### The Goldilocks zone

From these results we begin to see that recently isolated populations that experienced a strong bottleneck must satisfy particular conditions so that an allele at low frequency may rise quickly to fixation resulting in something resembling a classic selective sweep (Figure 1). First, the time since the colonization event and start of selection must be greater than the time to fixation, roughly ln(1/2*ƒ*)/*s* generations. On the other hand, this time should not be so long that genetic drift has erased the signal of the sweep, which occurs on the scale of *N*_*I*_ generations. Second, since the influx of new mutation is very small, adaptation likely comes from standing variation, and for it to be likely that the sweep is hard, *K* should be close to 1. These bounds delineate the Goldilocks zone in Figure 1. In particular, selection must be strong or else it will be overcome by drift, i.e. 2*s*_*Aa*_*N*_*I*_ must be greater than 1 (in the simulations, *s*_*Aa*_ = 0.05, as seen in Table 2 and Figure 4). On the other hand, a higher starting frequency can help ensure that an allele has a better chance of fixing, but this is likely to result in a haplotype signature that cannot be differentiated from a neutral background, as discussed next. Next we show that these theoretical considerations have the expected consequences on the practical ability to detect sweeps.

### Haplotype structure around the selected site

Whether or not an allele can fix is only half of the story when determining if detecting selection is possible. Genetic variation in the surrounding genomic region must also look different in some measurable way. Most tests for selection try to differentiate outlier regions of the genome from a neutral genetic background, so selection must impact the haplotype structure around the beneficial site. The lower the starting frequency of the beneficial allele and the higher the selection coefficient, the longer the associated haplotype structure is expected to be (Maynard Smith & Haigh 1974, Kaplan *et al*. 1989). To examine haplotype structure we look at windows of increasing length centered on the beneficial mutation and count the mean number of haplotypes in each window (Figure 5A-F).

**Fig. 5.**
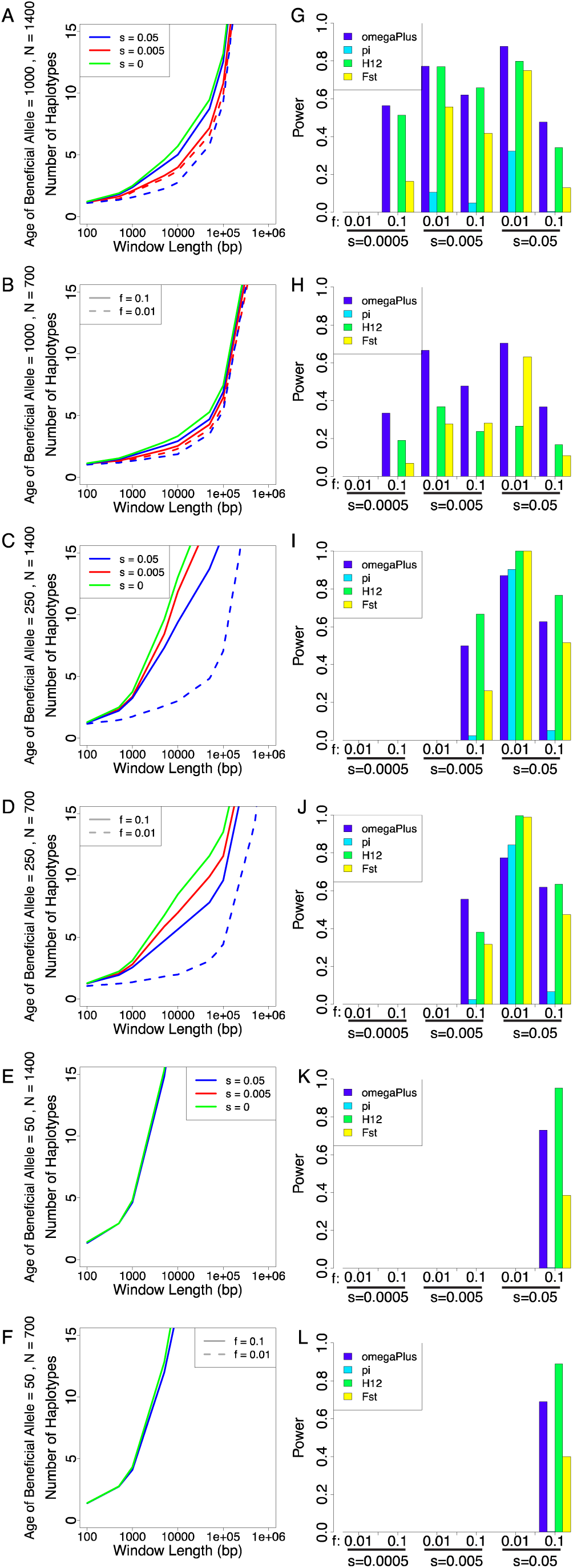
Extent of haplotype elongation around selected site and power of statistics. The age of the beneficial allele is given in generations, and *N* refers to size of the isolated population, *N*_I_ from Figure 2. Models with less than 20 replicates after filtering for *ƒ*_final_ ≥ 0.5 and at least 100 segregating sites are not shown in plots. A-F) Number of unique haplotypes in a window of fixed length surrounding the beneficial allele. The dotted lines represent a starting frequency of 0.01 and the solid lines represent a starting frequency of 0.1. G-L) Power for each of the 4 statistics, defned as the proportion of replicates, after filtering, whose maximum (or minimum for pi) value is greater than (less than) a threshold value, determined as the 99^th^ (1^st^) percentile of the corresponding neutral model.

In models where fixation is likely, we see the greatest effect on haplotype length from *ƒ* = 0.01 at the strongest selection coefficient after 250 generations have passed (Figure 5C-D, dashed blue line). From Table 3, the expected time for this model to reach a frequency of 0.5 is 78 generations, therefore the expected time to reach fixation is roughly twice this number, or 156 generations. For *ƒ* = 0.1, the allele has also fixed at this time point (estimated *T*_fix_ = 64 generations), but it does not produce the same elongated haplotype as the model with lower starting frequency. We see almost no difference in the haplotype structure at 50 generations, when it is expected to be first nearing fixation (Figure 5E-F). This is what we expect if we consider that there are estimated to be around 7-14 haplotypes that fix under this selection scenario (Table 2), i.e. the sweep is too soft (Przeworski *et al*. 2005).

After 1000 generations, while still producing the most severe impact on haplotype length, the signature of elongation for *ƒ* = 0.01 and *s* = 0.05 is more similar to the neutral model (Figure 5A-B). Closer inspection of Figure 5 reveals that this difference comes from the length of the neutral haplotype at the two different time points. At 250 generations, the allele has already fixed and eliminated variation that did not recombine onto the sweeping haplotype, creating a long block of shared genetic variation. Between 250 and 1000 generations, drift in the small population causes the haplotypes to become longer in the neutral model, which should give us less power to detect a selective sweep (and it does, see Figure 5G-K).

Weaker selection (*s*_*Aa*_ = 0.005), while still ultimately effective, does not show a strong haplotype signal (red lines in Figure 5A-B). This is because the sweep takes much longer to occur (Figure 4), leaving more time for the sweeping haplotype to recombine with others.

Another noticeable effect on haplotype length can be seen from the size of the bottleneck. Haplotypes for all models are slightly longer for the more severe reduction where *N*_*I*_=700 (*n* = 0.01). This closes the gap between the amount of genetic variation that is lost due to the sweep and that due to the bottleneck, making it more difficult to distinguish a sweep. This is because drift in the smaller population removes variation, producing longer haplotypes on average for all scenarios.

By considering haplotype length around the selected site one model stands out as showing the most distinct sweep-like signature: strong selection (*s*_*Aa*_ = 0.05) on a lower starting frequency (*ƒ* = 0.01). This combination is closest to a hard sweep from a de novo mutation, with only 1.4 alleles expected to fix in the largest population (Table 3). Of the three time points that were simulated, sampling when the allele is 250 generations old produces the most pronounced effect on the haplotype structure around the target site. In this scenario, the beneficial allele is old enough to have fixed, but not so old that drift has erased the signature of selection. This combination of parameters therefore lies in the Goldilocks zone, which is conceptually illustrated in Figure 1. Above we discussed that in order to observe selection, the time to fixation must be larger than ln(1/2*ƒ*)/*s*, and it is known that diversity within the target region will equilibrate after *N*_I_ generations (Przeworski 2002). If the expected number of alleles to fix, *K* = 2*s*_*Aa*_*N*_I_*ƒ*, is too large, and initial diversity is sufficient they are each on different haplotypes, the sweep will be too soft to detect a change in diversity around the selected site, and if it is too small it will not have a high enough chance of fixation. The values of *K* that we find to be ideal with the parameters of our demographic values surrounding our Goldilocks model are around 1 (0.7 or 1.4, Table 3, Figure 1).

### Performance of statistical tests in detecting a selective sweep

Next, we examined the power of four statistics to detect the signature of selection when a sweep occurs in each of our models. All four have the most power to detect a sweep in the Goldilocks zone described above, which leaves the strongest haplotype signature (Figure 5I-J).

Testing for reduced diversity (π) is generally underpowered in most models because there is not enough genetic variation in the population after the bottleneck, making it difficult to distinguish between the neutral and the selection models. Since π does not rely on differentiation, like *F*_ST_, or a specific pattern in the haplotypes, like omegaPlus and H12, it lacks power when genetic variation has been reduced globally due to a bottleneck.

The two linkage-based estimators we examined, omegaPlus and H12, perform similarly for all models, except that OmegaPlus has about twice the power of H12 when the beneficial allele is old (1000 generations) and the bottleneck is more severe, *N*_I_ = 700 (Figure 5H). Since H12 counts haplotypes in a fixed window of SNPs, it will have the most power to detect a sweep when the sample size is very large (here we only sample 50 diploid individuals.) A more severe bottleneck will result in fewer haplotypes fixing from selection (Table 2) and has a similar effect to reducing the sample size, or the number of haplotypes, for the whole region. This could make H12 a tricky statistic to use in conjunction with populations that are expected to have a severe reduction in size due to a recent colonization event.

One instance when H12 seems to do better than OmegaPlus is when the age of the beneficial allele is 50 generations (Figure 5K-L). In this case a high number of haplotypes are expected to have fixed (Table 2), within about the last 10 generations (Table 3). This increased diversity in haplotypes makes H12 more sensitive, because there is more room for a difference in the frequencies of haplotypes between the selected and neutral models. OmegaPlus relies on the sweep-like pattern of LD around a beneficial allele, and with increased SNP diversity, that pattern is weakened.

### Application to real examples

The different scenarios discussed here can be used to gauge the likelihood of detecting selection in real-life populations when some basic parameters about the founding event can be estimated. For the island mice that differ in their level of sperm competition, the population size on Rat Island is estimated to be around 772 individuals, close to our simulated model of *N*_I_ = 700. Whitlock Island has an estimated 111 individuals, most closely resembling the model with *N*_I_ = 70.

Our simulations show that with the severity of the bottleneck on Whitlock Island it is unlikely that alleles have fixed due to selection or if they have, that we can detect them. Any alleles that may have fixed, e.g. *s*_*Aa*_ = 0.05, *ƒ* = 0.01 for this value of *N*_I_ (Table 2) would have done so very quickly (within 64 generations, Table 3). Given that Whitlock was colonized more than *N*_I_ generations ago (estimated colonization time is 800 generations ago), no signature of a sweep would remain – genetic drift due to the extremely small population size eliminates variation very quickly. Indeed, many of the simulations with the most severe bottleneck were eliminated from further analysis because they didn’t have at least 100 segregating sites in the 1Mb region that was simulated.

For Rat Island the situation is less grim. Given its more recent colonization (within 300 generations), if the selection scenario lies in the Goldilocks zone, say, *s*_*Aa*_ at least 0.05 from *ƒ* = 0.01 then the beneficial allele will have time to fix (Table 3) and the signal should not have been completely eliminated by subsequent drift, since *N*_I_ = 700. At high starting frequency, *ƒ* = 0.1, the beneficial allele is not expected to reduce genetic variation to an extent where it will be easily distinguishable from the neutral model (Figure 5C-D). But if the target allele was at low frequency after colonization and became fixed in the population, it should be possible to distinguish this region using one of the statistics examined here.

Another population of island mice that falls within the Goldilocks zone is the Gough Island mice in the South Atlantic. It is estimated that the mice arrived around 110 generations ago and established a population of approximately *N*_I_ = 950 (Gray *et al*. 2014). With these values, a variant with *s*_Aa_ = 0.05 and *ƒ* = 0.01 is expected to have about K = 1.0 alleles fix. From Table 3, the time it would take for these alleles to fix is around 150 generations. Therefore, if the ecologically interesting phenotypes arose from standing variation on a small proportion of haplotypes then we should be able to distinguish the sub-genomic regions that were responsible for selection. Gray *et al*. (2014) also estimated growth after the bottleneck, from *N*_e_ around 950 to present *N*_e_ around 20,000. This should not affect the ability to detect selection, since there will not be much new mutation given the short time period.

### The general prospectus

We now more briefly evaluate other well-known instances of evolution in isolated populations. In the Goldilocks zone, the expected number of sweeping haplotypes, *K*, is close to 1, and the number of generations since isolation, *t*, is between *T* = ln(1/2*ƒ*)/*s* and *N*_e_. Since *K* is equal to the number of beneficial alleles in the founding population (2*N*_I_ *ƒ*) multiplied by the selection coefficient (*s*), selection is only likely to be detected if *s* is close to 1/2*N*_I_*ƒ*. Furthermore, so that the sweep has had time to complete, *s* must be at least ln(1/2*ƒ*)/*t*. At given values of *N*_I_ and *t*, these put an upper bound on the initial frequency of the selected allele, and a corresponding lower bound on the strength of selection that is likely to be detectable (and then only if the corresponding frequency matches); these are shown for each scenario in Table 1. A few arbitrarily chosen combinations of initial frequency and selection coefficient are also shown.

Table 1 shows, unsuprisingly, that strong selection is most likely to be detectable, if the initial frequency of the selected allele is appropriate. Large, old isolated populations (e.g. natural guppy population isolates, Endler 1980) should allow and retain signals of much weaker selective sweeps for much longer, but only if the signals have not been erased by drift (*t<N*_e_), and may be limited by the amount of founding genetic diversity.

Some have fairly narrow requirements: for instance, experimental guppy introductions described in Endler (1980) were resampled after 15 generations, and so selection must be fairly strong to fix in this short a time. However, since the initial population size was large, the corresponding initial frequency (*ƒ*) must be small, which forces *s* to be still larger; the smallest value of *s* that allows both *K* = 1 and *T* < *t* is s = 0.31. This assumes the post-introduction collapse to the long-term *N*_e_ of 30 was not immediate; if this happened immediately, the appropriate *N*_I_ would be 30, not 200.

## Conclusions

Our simulations highlight how tough it can be to identify sites experiencing selection from whole genome scans in isolated populations that have experienced a recent bottleneck. These populations can provide a rich opportunity to study evolution in action, but the timeframe for selection or the extent of genetic drift due to reduced population size can hamper the ability to identify the genetic basis of adaptation with genomic outlier scans.

Many of the tools that are used in genomic outlier scans look for the reduction in diversity accompanying a classic hard sweep. However, this mode of selection may not be realistic for complex adaptive phenotypes, such as the evolution of body size or mating dynamics. An important focus of future research could be to see if our conclusions hold under a polygenic model. Also, initial genetic adaptation in small populations will likely occur from standing genetic variation. Soft sweeps from standing variation leave a weak signal in the genome, making it difficult to uncover selected sites from genomic data. In our simulations of this scenario, statistics like H12 have more power to find selected sites if the sample size is large and the beneficial allele has multiple origins. However, with a small population size, the window of time where H12 has an advantage is small because of the diversity-reducing effects of genetic drift. Since the action and mode of selection is likely not ubiquitous across the genome, in the future the best exploratory genomic approach should be one that encompasses several statistics that have strengths in identifying a range of adaptive signals.

The conclusions are not entirely pessimistic. First, we have not considered the increased power possible using replicate populations, which is common in experimental situations. For this to be successful the genetic basis must be shared among the replicates; this may not be the case if there are many possible adaptations, even if within each population stochasticity only chooses one. Second, although some of the parameters are not within the researcher’s control, others are, either by direct manipulation or by choosing appropriate study systems.

Although we only simulated 1 megabase of genome in 50 diploids, the implications are clear for whole-genome studies. Larger sample sizes should generally give more power, especially when it comes to haplotype-based statistics, but will not allow inference much more outside of the Goldilocks zone, since theoretical constraints still hold. Longer sequences will naturally make the identifying selected sites harder, unless the number of selected sites scales concordantly.

A final point that we have not considered is the genomic resolution of the outlier scan, when successful. In general, the resolution should be better with larger populations and weaker selection (Przeworski 2002), although the increased stochasticity accompanying weaker selection may be a problem. The guidelines we give here should be useful for back-of-the-envelope calculations, but are not a substitute for detailed power simulations tailored to particular situations.

## Acknowledgements

Funding for this study was provided by National Science Foundation grant #1146525 (MDD) and USC startup funds (PR). Emily Kopania helped code application of the H12 statistic. Lorraine Provencio assisted with Table 1.

## Data Accessibility

Scripts used in producing and analyzing the simulations are available upon request.

## References

Akey JM (2009) Constructing genomic maps of positive selection in humans: where do we go from here? Genome Research, 19, 711–722.

Alachiotis N, Stamatakis A, Pavlidis P (2012) OmegaPlus: a scalable tool for rapid detection of selective sweeps in whole-genome datasets. Bioinformatics, 28, 2274–2275.

Berry RJ (1996) Small Mammal Differentiation on Islands. Philosophical Transactions of the Royal Society B: Biological Sciences, 351, 753–764.

Burke MK, Dunham JP, Shahrestani P et al. (2010) Genome-wide analysis of a long-term evolution experiment with Drosophila. Nature, 467, 587–590.

Coop G, Ralph P (2012) Patterns of neutral diversity under general models of selective sweeps. Genetics, 192, 205–224.

Cuthbert R, Hilton G (2004) Introduced house mice Mus musculus: a significant predator of threatened and endemic birds on Gough Island, South Atlantic Ocean? Biological Conservation, 117, 483–489.

Endler JA (1980) Natural Selection on Color Patterns in Poecilia reticulata. Evolution, 34, 76.

Firman RC, Klemme I, Simmons LW (2013) Strategic adjustments in sperm production within and between two island populations of house mice. Evolution, 67, 3061–3070.

Firman RC, Simmons LW (2008) The frequency of multiple paternity predicts variation in testes size among island populations of house mice. Journal of Evolutionary Biology, 21, 1524–1533.

Firman RC, Simmons LW (2011) Experimental evolution of sperm competitiveness in a mammal. BMC Evolutionary Biology, 11, 19.

Fisher R (1918) The correlation between relatives on the supposition of Mendelian Inheritance. Transactions of the Royal Society of Edinburgh, 52, 399–433.

Fraser BA, Künstner A, Reznick DN, Dreyer C, Weigel D (2015) Population genomics of natural and experimental populations of guppies (Poecilia reticulata). Molecular Ecology, 24, 389–408.

Garud NR, Messer PW, Buzbas EO, Petrov DA (2015) Recent Selective Sweeps in North American Drosophila melanogaster Show Signatures of Soft Sweeps. PLoS Genetics, 11, 1–32.

Gill AE (1977) Maintenance of polymorphism in an island population of the California vole, Microtus californicus. Evolution, 31, 512–525.

Gray MM, Wegmann D, Haasl RJ et al. (2014) Demographic History of a Recent Invasion of House Mice on the Isolated Island of Gough. Molecular Ecology, 23, 1923–1939.

Haldane J (1924) A mathematical theory of natural and artificial selection. Transactions of the Cambridge Philosophical Society, 23, 19–41.

Hermisson J, Pennings PS (2005) Soft sweeps: molecular population genetics of adaptation from standing genetic variation. Genetics, 169, 2335–2352.

Holland B, Rice WR (1999) Experimental removal of sexual selection reverses intersexual antagonistic coevolution and removes a reproductive load. Proceedings of the National Academy of Sciences, 96, 5083–5088.

Kaplan NL, Hudson RR, Langley CH (1989) The “hitchhiking effect” revisited. Genetics, 123, 887–899.

Keller SR, Taylor DR (2008) History, chance and adaptation during biological invasion: Separating stochastic phenotypic evolution from response to selection. Ecology Letters, 11, 852– 866.

Kim YH, Nielsen R (2004) Linkage disequilibrium as a signature of selective sweeps. Genetics, 167, 1513–1524.

Kolbe JJ, Leal M, Schoener TW, Spiller DA, Losos JB (2012) Founder effects persist despite adaptive differentiation: a field experiment with lizards. Science (New York, N.Y.), 335, 1086–9.

Ledevin R, Chevret P, Ganem G et al. (2016) Phylogeny and adaptation shape the teeth of insular mice group. Proceedings of the Royal Society B: Biological Sciences, 283, 20152820.

Lescak EA, Bassham SL, Catchen J et al. (2015) Evolution of stickleback in 50 years on earthquake-uplifted islands. Proceedings of the National Academy of Sciences, 112, 201512020.

Losos JB, Ricklefs RE (2009) Adaptation and diversification on islands. Nature, 457, 830–836.

Losos JB, Schoener TW, Warheit KI, Creer D (2001) Experimental studies of adaptive differentiation in Bahamian Anolis lizards. Genetica, 112, 399–415.

Martinkova N, Barnett R, Cucchi T et al. (2013) Divergent evolutionary processes associated with colonization of offshore islands. Molecular Ecology, 22, 5205–5220.

Maynard Smith JM, Haigh J, Smith JM (1974) The hitch-hiking effect of a favourable gene. Genetical Research, 23, 23–35.

McVean G (2007) The structure of linkage disequilibrium around a selective sweep. Genetics, 175, 1395–1406.

Patton JL, Yang SY, Myers P (1975) Genetic and morphologic divergence among introduced rat populations (Rattus rattus) of the Galápagos Archipelago, Ecuador. Systematic Biology, 24, 296– 310.

Pergams ORW, Byrn D, Lee KLY, Jackson R (2015) Rapid morphological change in black rats (Rattus rattus) after an island introduction. PeerJ, 3, e812.

Pergams ORW, Lawler JJ (2009) Recent and widespread rapid morphological change in rodents. PLoS ONE, 4, e6452.

Poh YP, Domingues VS, Hoekstra HE, Jensen JD (2014) On the Prospect of Identifying Adaptive Loci in Recently Bottlenecked Populations. PLoS ONE, 9, e110579.

Przeworski M (2002) The signature of positive selection at randomly chosen loci. Genetics, 160, 1179–1189.

Przeworski M, Coop G, Wall JD (2005) The Signature of Positive Selection on Standing Genetic Variation. Evolution, 59, 2312–2323.

Reznick DN, Ghalambor CK (2001) The population ecology of contemporary adaptations: What empirical studies reveal about the conditions that promote adaptive evolution. Genetica, 112, 183–198.

Reznick DN, Bryga H (1987) Life-history evolution in guppies (Poecilia reticulata): 1. Phenotypic and genetic changes in an introduction experiment. Evolution, 41, 1370–1385.

Reznick D, Endler JA (1982) The Impact of Predation on Life History Evolution in Trinidadian Guppies (Poecilia reticulata). Evolution, 36, 160–177.

Rowe-Rowe D, Crafford J (1992) Density, body size, and reproduction of feral house mice on Gough Island. South African Journal of Zoology, 27, 1–5.

Swallow JG, Koteja P, Carter PA, Garland T (1999) Artificial selection for increased wheel-running activity in house mice results in decreased body mass at maturity. Journal of Experimental Biology, 202, 2513–2520.

Wright S (1950) Genetical structure of populations. Nature, 166, 247–249.

Wright S, Dobzhansky T (1946) Genetics of natural populations; experimental reproduction of some of the changes caused by natural selection in certain populations of Drosophila pseudoobscura. Genetics, 31, 125–156.

